# Combination attenuation offers strategy for live-attenuated coronavirus vaccines

**DOI:** 10.1101/309591

**Authors:** Vineet D. Menachery, Lisa E. Gralinski, Hugh D. Mitchell, Kenneth H. Dinnon, Sarah R. Leist, Boyd L. Yount, Eileen T. McAnarney, Rachel L. Graham, Katrina M. Waters, Ralph S. Baric

## Abstract

With an ongoing threat posed by circulating zoonotic strains, new strategies are required to prepare for the next emergent coronavirus (CoV). Previously, groups had targeted conserved coronavirus proteins as a strategy to generate live-attenuated vaccine strains against current and future CoVs. With this in mind, we explored whether manipulation of CoV NSP16, a conserved 2’O methyltransferase (MTase), could provide a broad attenuation platform against future emergent strains. Using the SARS-CoV mouse model, a NSP16 mutant vaccine was evaluated for protection from heterologous challenge, efficacy in the aging host, and potential for reversion to pathogenesis. Despite some success, concerns for virulence in the aged and potential for reversion makes targeting NSP16 alone an untenable approach. However, combining a 2’O MTase mutation with a previously described CoV fidelity mutant produced a vaccine strain capable of protection from heterologous virus challenge, efficacy in aged mice, and no evidence for reversion. Together, the results indicate that targeting the CoV 2’O MTase in parallel with other conserved attenuating mutations may provide a platform strategy for rapidly generating live-attenuated coronavirus vaccines.

**Significance:** Emergent coronaviruses remain a significant threat to global public health and rapid response vaccine platforms are needed to stem future outbreaks. However, failure of many previous CoV vaccine formulations has clearly highlighted the need to test efficacy under different conditions and especially in vulnerable populations like the aged and immune-compromised. This study illustrates that despite success in young models, the NSP16 mutant carries too much risk for pathogenesis and reversion in vulnerable models to be used as a stand-alone vaccine strategy. Importantly, the NSP16 mutation can be paired with other attenuating approaches to provide robust protection from heterologous challenge and in vulnerable populations. Coupled with increased safety and reduced pathogenesis, the study highlights the potential for NSP16 attenuation as a major component of future live-attenuated coronavirus vaccines.

## Introduction

The emergence of severe acute respiratory syndrome coronavirus (SARS-CoV) at the beginning of the 21st century signaled a new era for emergent viral disease (1). Since then, dozens of viruses have emerged from animal populations to produce significant outbreaks in humans and transfer of zoonotic pathogens remains a major threat to global public health (2). A decade after SARS-CoV, infections with Middle East respiratory syndrome coronavirus (MERS-CoV) continue with periodic reintroductions still occurring six years after its initial discovery (3). Coupled with recently discovered SARS-and MERS-like coronaviruses circulating in animal populations, the threat of a coronavirus fueled outbreak remains far from remote (4). With this in mind, strategies must be developed to rapidly respond to a potential CoV emergence or reemergence event.

While the SARS-and to a lesser extent the MERS-CoV outbreaks have been mostly contained, this limitation has been primarily due to effective public health measures and the delayed transmissibility of the viruses until after symptomatic disease (5). Standard antiviral treatments with type I IFN or traditional nucleoside analogs like ribavirin have shown minimal success against either epidemic strain (6). While several antibodies have been designed against SARS-and MERS-CoV, their efficacy has been found to be strain specific and may offer minimal protection against heterologous or unrelated coronaviruses, especially under therapeutic conditions (4, 7). Other drugs, targeting conserved CoV activities, have had either marginal efficacy *in vivo* or offer only a small window for treatment during early infection (8, 9). Similar to other viral infections, the most effective treatment for coronaviruses is prevention via vaccination.

In the context of both the SARS-and MERS-CoV outbreaks, a wealth of vaccine strategies have been developed and examined (10, 11). Many have utilized sub-unit based approaches; others have used virus like particles or other vector systems to induce immunity. While less desirable for both commercial and safety reasons, a number of live attenuated strains have also been developed targeting both conserved CoV elements or specific approaches that may be strain or group dependent (12, 13). For many of these vaccine approaches, protection from homologous challenge was noted following *in vivo* vaccination of young animals (10, 11). However, the failure of non-infectious vaccines in regards to heterologous challenge and in aged mice signaled a major issue in their use in vulnerable populations most impacted by coronavirus disease (14, 15). While some success was observed in aged mice and with heterologous challenge, live-attenuated coronavirus vaccines have significant concerns for reversion and pathogenesis (16, 17). Together, the results indicate the need for an improved platform for coronavirus vaccines and is a major reason MERS-CoV has been included as a target by the Coalition for Epidemic Preparedness Innovations (CEPI).

In this manuscript, we expand upon a live-attenuated vaccine approach based on mutation of coronavirus NSP16, a 2’O methyl-transferase. Previous work in murine hepatitis virus, SARS-CoV, and MERS-CoV have found that disruption of NSP16 activity rendered an attenuated strain sensitive to the activity of interferon stimulated IFIT1 (18-20). Work by our group went on to show that NSP16 mutants provided protection from lethal challenge by both SARS-CoV and MERS-CoV in young animals (19, 20). In this work, we set forth to evaluate NSP16 attenuation as a platform strategy for live-attenuated coronavirus vaccines. Using models that explore heterologous challenge, efficacy in aging, and potential for reversion, we found that the NSP16 mutant alone was not sufficiently attenuated to be used universally. However, the broad conservation of NSP16 coupled with robust viral yields permitted its use in combination with another attenuating mutation in NSP14, exonuclease (ExoN) activity. Targeting both conserved coronavirus activities produced a stable, attenuated virus capable of protection from heterologous challenge, efficacy in aged mice, and absence of reversion in immunocompromised models. Together, the results indicate that targeting coronavirus NSP16 may be a critical component of a future live attenuated coronavirus vaccine approach.

## Results

Previous live-attenuated coronavirus platforms had taken advantage of the need and conservation of NSP14 exonuclease (ExoN) and the coronavirus envelope (E) proteins (16, 21). Similar to ExoN, the NSP16 2’O MTase has significant homology across the coronavirus family, more than CoV envelope proteins (**Fig. 1A**). Importantly, the critical KDKE enzymatic motif is highly conserved allowing predictable disruption of the 2’O MTase function (2). Unlike ExoN and E viral mutants, dNSP16 viruses have already been shown to attenuate infection of several coronavirus strains including MERS-CoV (20, 22). In addition, for both SARS-and MERS-CoV, the NSP16 mutant viruses induce robust protection following lethal homologous challenge *in vivo* (19, 20). However, despite proof of concept studies, several questions about the efficacy of the NSP16 vaccine remain. With the absence of replication attenuation, it is especially important to consider the NSP16 vaccine platform in the context of the overall host response. In addition, evaluation of heterologous challenge, aged models of disease, and the potential for reversion represent critical factors that must be examined prior to further pursuit as a live-attenuated vaccine platform.

**Figure 1.**
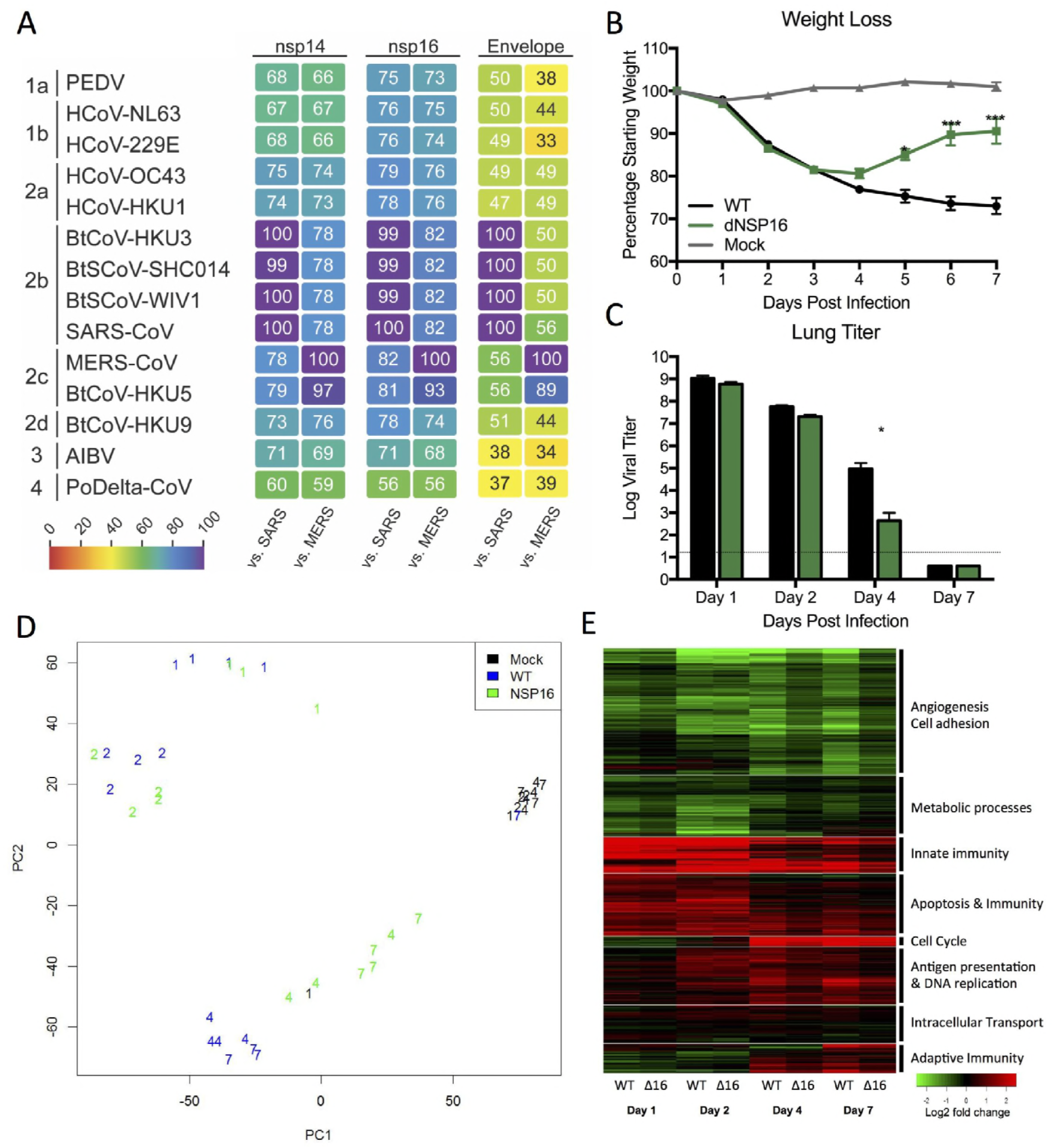
Host response to SARS-CoV dNSP16 CoV mirrors wild-type at early times. A) NSP14 exonuclease (ExoN), NSP16 (2’O MTase), and envelope (E) protein sequences of the indicated viruses were aligned and sequence identities were extracted from the alignments. A heatmap of sequence identity using SARS-CoV Urbani or MERS-CoV EMC as the reference sequence was constructed using Evolview (www.evolgenius.info/evolview). The heatmap was further rendered and edited in Adobe Illustrator CC 2017. B-C) Twenty-week old C57BL/6 mice challenged with mock (PBS, gray), SARS-CoV MA15 (WT, black), or SARS-CoV dNSP16 (green) at 10^5^ PFU were examined for B) weight loss (n > 5 for WT and dNSP16 groups) and C) lung viral titer (n = 5 per group). D) PCA analysis of total RNA expression at individual times for WT (blue), dNSP16 (green), and mock (black). Numerical marking (1, 2, 4, & 7) indicate time point of sample. E) Differentially expressed genes relative to mock were used to generate clustered expression heat maps comparing expression in WT and dNSP16 following *in vivo* infection across the time course. P-values based on Student T-test and are marked as indicated: * < 0.05 *** < 0.001.

### Early host response to SARS-CoV NSP16 mutant equivalent to wild-type infection

Unlike the ExoN and E protein mutants, dNSP16 coronaviruses demonstrate no significant attenuation in terms of viral replication *in vitro* except in the context of type I IFN pretreatment (9). Similarly, *in vivo* characterizations found no deficit in viral replication at early times (days 1 and 2), but a marked attenuation at late times post infection (19, 20). These results suggest that the early portions of dNSP16 infection are equivalent to wild-type virus infection. To further explore this question, we utilized a systems biology approach to compare wild-type and dNSP16 mutant in 20-week old C57BL/6 mice. In this model, mice infected with the mouse-adapted SARS-CoV induce severe weight loss, fail to recover, and several succumb to infection (**Fig. 1B**). Notably, while the SARS NSP16 mutant induces weight loss, mice recover from infection diverging from wild-type infection 3 days post infection. Examining viral replication, the NSP16 mutant has no discernable attenuation at days 1 or 2 (**Fig. 1C**); however, at day 4, a 100-fold reduction in replication is observed in the lungs of dNSP16 infected mice that corresponds with the kinetics of interferon stimulated gene (ISG) expression. Importantly, the global host expression response models the same trend. PCA plots of total RNA expression finds that wild-type and mutant virus profiles cluster together at days 1 and 2 (**Fig. 1D**). At later times, similar to weight loss and viral replication, dNSP16 diverges from WT, suggesting that only at late times do the host responses change following infection with the 2’O MTase mutant. Network analysis found similar results, with overlapping signatures at early time points (**Fig. 1E**). In contrast, late RNA expression demonstrated waning host responses consistent with reduced viral loads. In this context, the observation could be seen as a positive signal for inducing robust immune responses for vaccine responses. Conversely, the overlap with WT host responses may indicate a potential risk for inducing significant damage prior to attenuation of infection at late times.

### NSP16 vaccine protects from heterologous challenge

With the continued circulation of SARS-and MERS-like viruses in animal populations worldwide (4), vaccine approaches must consider exposure to and protection from related, heterologous coronavirus strains. Utilizing a chimeric mouse adapted SARS-CoV strain incorporating the spike of SHC014-CoV (7), an antigenically distant group 2B coronavirus circulating in Chinese horseshoe bat populations, we evaluated the efficacy of the SARS-CoV dNSP16 in protecting from heterologous CoV challenge. BALB/c mice were vaccinated with 10^5^ PFU of SARS dNSP16, monitored for 28 days, and subsequently challenged with the chimeric strain (SHC014-MA15). Our results indicate that the attenuated 2’O MTase SARS-CoV mutant is capable of inducing robust protection from the zoonotic group 2B spike virus in terms of disease. Weight loss data, viral lung titer, and hemorrhage score indicated little to no disease in mice vaccinated with dNSP16 compared to mock vaccinated mice (**Fig. 2A-C**). Importantly, while not equivalent to neutralization of wild-type SARS-CoV, sera from dNSP16-vaccinated mice were able to effectively block the SHC014 chimeric virus, likely contributing to protection following vaccination (**Fig. 2D**). Together, the data indicate that a live-attenuated NSP16 vaccine has sufficient capacity to protect from heterologous CoV challenge in young mice.

**Figure 2.**
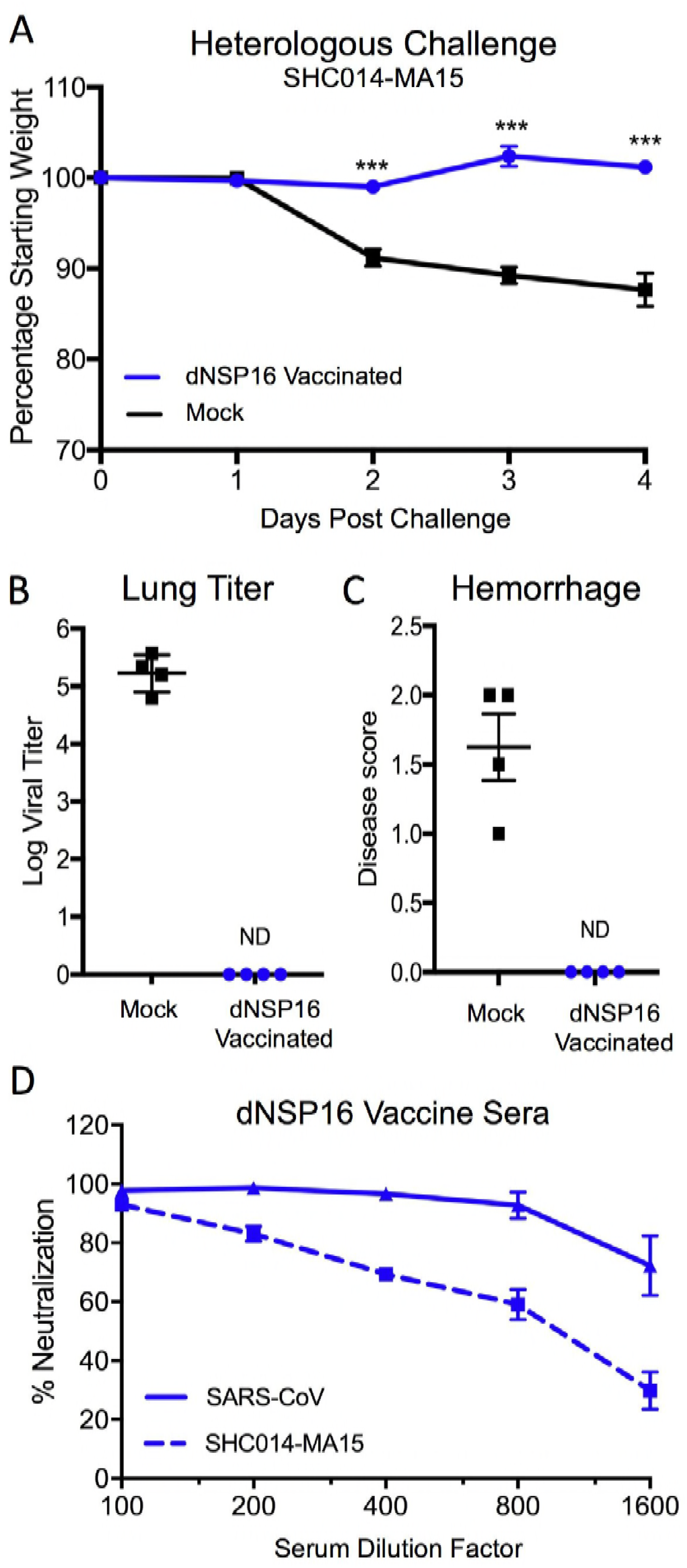
NSP16 vaccine protects from heterologous challenge. A-C) Ten-week old BALB/C mice were vaccinated with 10^5^ PFU of SARS-CoV dNSP16 (blue) or mock (PBS), monitored for 28 days, and subsequently challenged with 10^5^ PFU of heterologous SARS-CoV expressing SHC014 spike (SHC014-MA15). Mice were examined over a four-day time course for A) weight loss, B) day 4 lung virus titer, and C) day 4 lung hemorrhage. D) Plaque reduction neutralization titers of WT SARS-CoV (solid line) or heterologous SHC014-MA15 (hashed line) from sera of dNSP16 vaccinated. P-values based on Student T-test and are marked as indicated: *** < 0.001.

### Low dose dNSP16 vaccination provides protection to aged mice

With high rates of mortality, aged populations represent a vulnerable group in need of effective vaccines to coronaviruses (23). However, previous work has shown that SARS-CoV vaccines have had mixed success in aged populations (14, 24). To explore efficacy of dNSP16 in a senescent model, we examined infection of 12-month old BALB/c mice which recapitulates the age-dependent susceptibility observed in humans. Following infection with 10^5^ PFU of wild-type or dNSP16, the 2’O MTase mutant was equal to WT SARS-CoV infection in terms of weight loss and lethality (**Fig. 3A & B**). While this dose protected young mice from heterologous challenge, significant pathogenesis occurred in the aged mice despite attenuation in viral replication at day 4 (**Fig. 3C**). Together, the data indicates that despite replication attenuation, the dNSP16 mutant harbors pathogenic potential at high doses in aged mice.

**Figure 3.**
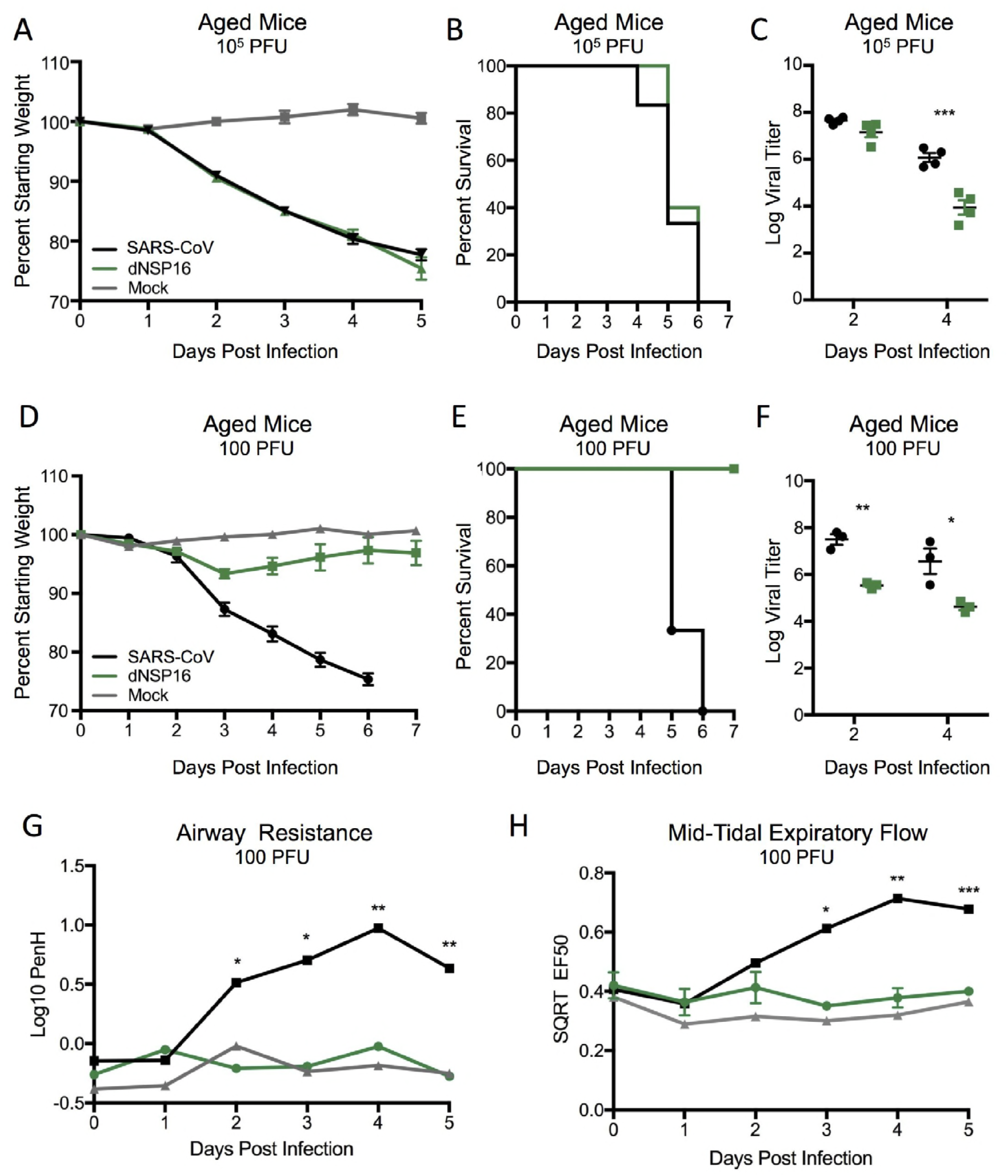
SARS-CoV dNSP16 infection risks virulence in aged mice. A-C) Twelve-month old BALB/c mice were challenged with 10^5^ PFU of WT SARS-CoV MA15 (black), dNSP16 (green) or mock (PBS, grey) and examined for A) weight loss, B) survival, and C) lung viral titer over the time course. D-F) Twelve-month old BALB/c mice were challenged with 100 PFU of WT SARS-CoV MA15 (black), dNSP16 (green), or mock (PBS, gray) and examined for D) weight loss, E) survival, F) lung viral titer, G) airway resistance (PenH) and H) Mid-Tidal Expiratory Flow (EF50) as measured by whole body plethysmography (Buxco, DSI). P-values based on Student T-test and are marked as indicated: * < 0.05, ** < 0.01, *** < 0.001.

While capable of inducing lethal disease, the utilized dose of the dNSP16 mutant represents a significant increase over the LD_50_ of WT SARS-CoV in aged mice (5). Therefore, we challenged 12-month old BALB/c mice with 100 PFU of WT or dNSP16 and evaluated pathogenesis. Following low dose challenge, dNSP16 infected aged mice had reduced disease as compared to control in terms of weight loss and survival (**Fig. 3D & E**). The reduced pathogenesis corresponded to reduced viral replication at both days 2 and 4, recapitulating *in vivo* attenuation previously observed in young animals (**Fig. 3F**) (9). Notably, dNSP16 infection resulted in only marginal disease in terms of weight loss (<5%) and respiratory parameters, including airway resistance (PenH, **Fig. 3G**) and mid-tidal expiratory flow (EF50, **Fig. 3H**). To explore protection, 12-month old mice were vaccinated with 100 PFU of dNSP16 and subsequently challenged 28 days later with a lethal dose of wild-type SARS-CoV MA15 (10^5^ PFU). Both, in terms of weight loss and survival (**Fig. 4A & B**), vaccination with dNSP16 provided robust protection from lethal disease and pathogenesis; this protection corresponded with the absence of detectable viral replication at both days 2 and 4 (**Fig. 4I**). Together, the results indicate that while the dNSP16 mutant harbors some risk of pathogenicity at high doses, it can also provide aged mice with robust protection from lethal challenge.

**Figure 4.**
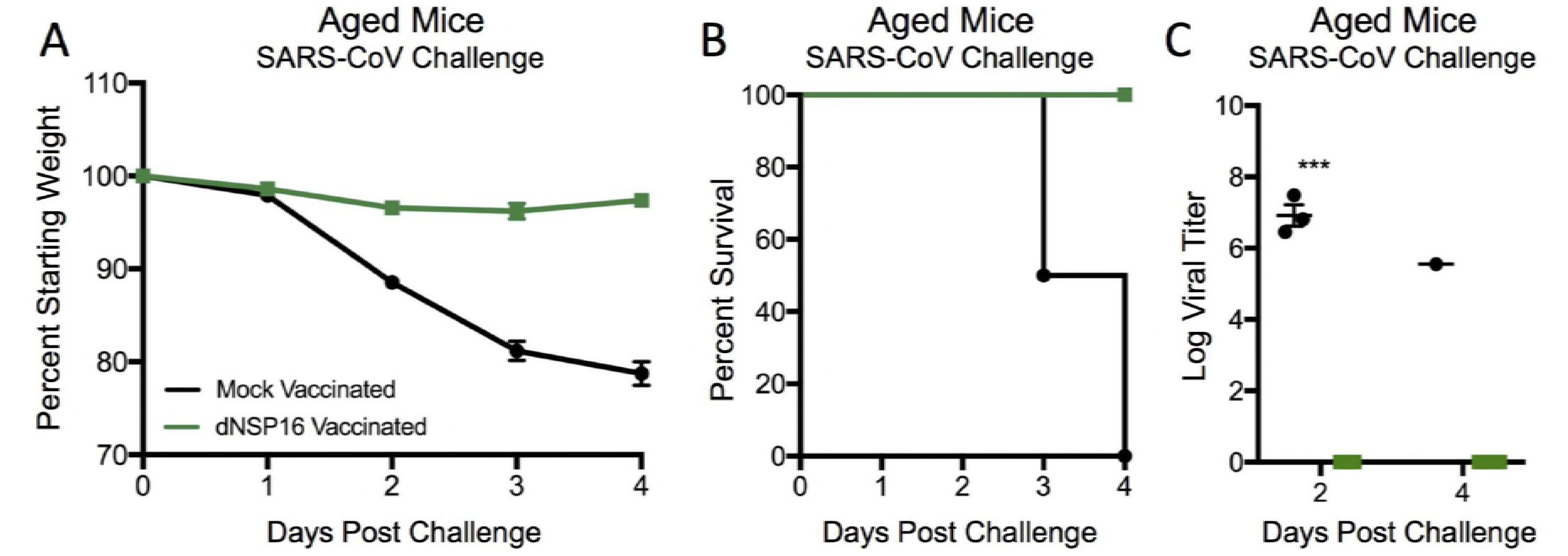
SARS-CoV dNSP16 vaccination protects aged mice. A-C) Twelve-month old BALB/c mice were vaccinated with 100 PFU of SARS-CoV dNSP16 or mock (PBS), monitored for 28 days, and subsequently challenged with 10^5^ PFU of WT SARS-CoV MA15. Mice were examined over a four day time course for A) weight loss, B) survival, and C) viral titer. P-values based on Student T-test and are marked as indicated: * < 0.05, ** < 0.01, *** < 0.001.

### NSP16 mutant reverts in immune-compromised model

Despite some important caveats, the dNSP16 mutant has demonstrated efficacy as a vaccine following both, heterologous challenge and in aged models of SARS-CoV disease. However, reversion and adaptation of a live attenuated strain represents a significant concern for its use in response to human epidemics. Previous studies with SARS-CoV found that mice lacking functional B and T-cells (RAG^-/-^) were unable to clear virus allowing continuous passage *in vivo (16)*; these studies seek to utilize this model to examine dNSP16 infection for the possibility of reversion or adaptations resulting in virulence. Following infection with the SARS dNSP16 mutant, RAG^-/-^ mice were observed for disease and lungs harvested 30 days post infection; individual lungs from 8 mice were subsequently homogenized, clarified, and used to inoculate 10-week old BALB/c mice for reversion to virulence. The data demonstrated that for 5 of the 8 *in vivo* passaged virus samples, continuous infection resulted in restored virulence including weight loss and lethality (**Fig. 5A & 5B**). Importantly, examination of input inoculum revealed that the three samples that had failed to induce weight loss also had no detectable virus (Fig. 5C). With this in mind, the input virus samples were amplified on Vero cells and the RNA sequence of NSP16 was examined for reversion. Surprisingly, the results found that all five virus positive samples retained the D130A mutation in NSP16 suggesting that reversion to virulence was driven by mutations at other locations in the SARS-CoV backbone. Overall, the results indicate that the constructed dNSP16 mutant is not currently a viable vaccine candidate due to reversion to virulence in immune-compromised populations.

**Figure 5.**
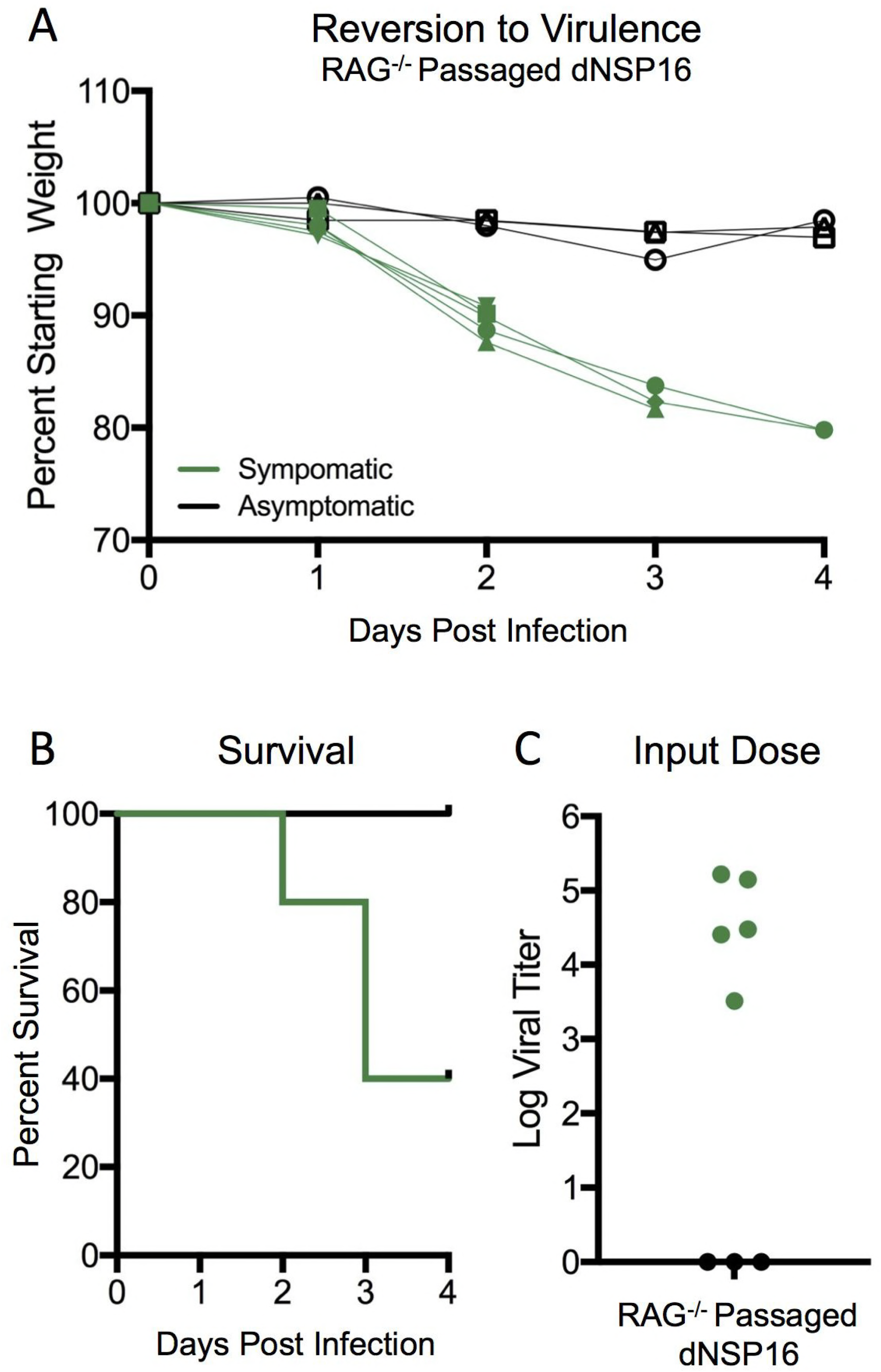
SARS-CoV dNSP16 can revert to virulence. A-C) Eight RAG^-/-^ mice were infected with 10^5^ PFU of SARS-CoV dNSP16, monitored, euthanized, and lung tissues harvested after 30 days. Clarified homogenates were then inoculated into 10-week old BALB/c mice and individually monitored for A) weight loss, B) survival, and C) input titer prior to infection. Five of eight RAG^-/-^ homogenates (green) produced significant weight loss, lethality, and had detectable input titers. Three RAG^-/-^ homogenates (black), had no disease and had no detectable virus.

### Viability of a multiple mechanism, live-attenuated vaccine

With concerns for reversion and potential virulence, a live-attenuated vaccine platform targeting only NSP16 activity should be limited to only dire circumstances. One possibility is to expand the NSP16 mutant beyond the original D130A mutations (2 nucleotides) to include other members of the KDKE motif (2). Another possibility was to leverage the NSP16 mutation in combination with other CoV live-attenuation vaccine platforms. Unlike strategies that target fidelity (ExoN) and the envelope protein, coronavirus NSP16 mutants have only minimal replication attenuation and demonstrate sensitivity to IFN stimulated genes after initial infection. With this in mind, we set forth to evaluate a SARS-CoV mutant lacking both NSP16 activity as well as CoV fidelity (dNSP16/ExoN). Compared to WT or the original dNSP16 mutant, the SARS dNSP16/ExoN virus had replication attenuation in its initial stock titers as well as in a multistep growth curve (**Fig. 6A**); however, the double mutant had equivalent replication to the SARS ExoN mutant indicating no additional replication attenuation *in vitro. In vivo,* the SARS dNSP16/ExoN mutant also had significant attenuation relative to WT control virus. Following infection of 10-week old BALB/c, dNSP16/ExoN produced no weight loss with significant changes compared to WT infection as early as day 2 post infection (**Fig. 6B**). Similarly, viral replication was significantly attenuated at both days 2 and 4 (**Fig. 6C**). Finally, no signs of hemorrhage were observed following infection with the double mutant (**Fig. 6D**). Together, the data is consistent with the level of attenuation observed for the original SARS-CoV ExoN mutant and suggested that the dNSP16/ExoN mutant may serve as a possible live attenuated vaccine.

**Figure 6.**
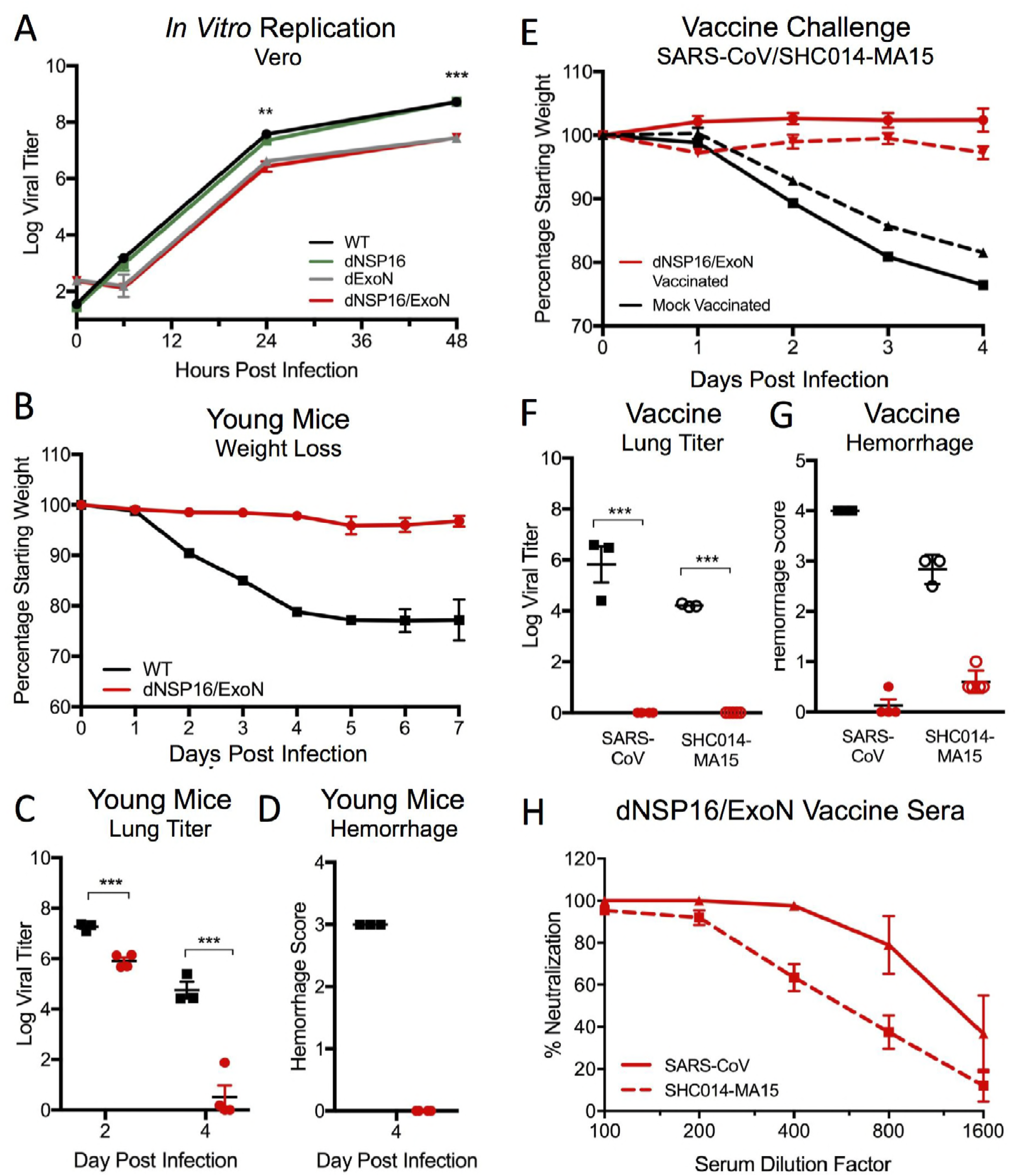
SARS-CoV dNSP16/ExoN is a viable vaccine and protects from homologous and heterologous challenge. A) Vero cells infected at a MOI 0.01 with WT SARS-CoV MA15 (black), dNSP16 (green), dExoN (grey), or dNSP16/ExoN (red). B-D) Ten-week old BALB/C mice were challenged with 10^4^ PFU of SARS-CoV MA15 (black) or dNSP16/ExoN (red) and examined for A) weight loss, B) lung viral titer, and C) lung hemorrhage. E-G) Ten-month old BALB/c mice were vaccinated with 10^4^ PFU of SARS-CoV dNSP16/ExoN (red) or mock (PBS, black), monitored for 28 days, and subsequently challenged with 10^5^ PFU of WT SARS-CoV MA15 (solid line) or SHC014-MA15 (dotted line). Mice were examined over a four day time course for E) weight loss, F) viral titer, and G) lung hemorrhage. H) Plaque reduction neutralization titers of WT SARS-CoV MA15 (solid) or heterologous SHC014-MA15 (dotted) from sera from dNSP16/ExoN vaccinated mice. P-values based on Student T-test and are marked as indicated: *** < 0.001.

With vaccination in mind, we explored the dNSP16/ExoN mutant in the context of homologous and heterologous SARS-CoV challenge. Young 10-week old BALB/c mice were vaccinated with dNSP16/ExoN or mock, monitored for 28 days, and subsequently challenged with homologous (SARS-MA15) or heterologous chimeric virus (SHC014-MA15). Following vaccination, both homologous and heterologous challenge induced significant weight loss in mock-vaccinated animals (**Fig. 6E**); in contrast, mice vaccinated with dNSP16/ExoN were protected. Similarly, dNSP16/ExoN vaccination resulted in complete ablation of viral titer at day 4 post infection following both homologous and heterologous challenge. In addition, hemorrhage scores indicated limited disease following vaccination with dNSP16/ExoN as compared to control (**Fig. 6F**). Importantly, sera from dNSP16/ExoN vaccinated mice was capable of neutralizing both WT SARS-CoV as well as chimeric spike virus (**Fig. 6H**). While not equivalent to neutralization induced by the dNSP16 mutant (**Fig. 2**), the results indicate that the double mutant vaccine provided robust protection from both homologous and heterologous challenge.

### Double mutant protects aged mice from lethal disease

Having demonstrated efficacy in young mice and against heterologous challenge, we next explored the SARS-CoV dNSP16/ExoN mutant in the context of an aging host. Twelve-month old BALB/c mice were challenged with 100 PFU of WT SARS-CoV (MA15) or the dNSP16/ExoN mutant. In terms of both weight loss (**Fig. 7A**) and lethality (**Fig. 7B**), the dNSP16/ExoN mutant was highly attenuated compared to the lethal WT infection. Similar to the young mouse challenge, the dNSP16/ExoN mutant had attenuated replication at days 2 and 4 post infection (**Fig. 7C**) and no signs of hemorrhage (**Fig. 7D**). Importantly, mice vaccinated with dNSP16/ExoN were completely protected from lethal SARS-CoV challenge. Both weight loss and survival showed complete protection in the vaccinated animals (**Fig. 7E**); in contrast, WT infected mice had significant weight loss and 2/3 succumbed to infection by day 4. Titers in the lung demonstrated no viral replication in vaccinated aged mice; in contrast, the lone surviving mock-vaccinated mice had significant titer within the lung (**Fig. 7F**). Together, the data demonstrated the utility and efficacy of the dNSP16/ExoN mutant as a live attenuated vaccine in aged models of disease.

**Figure 7.**
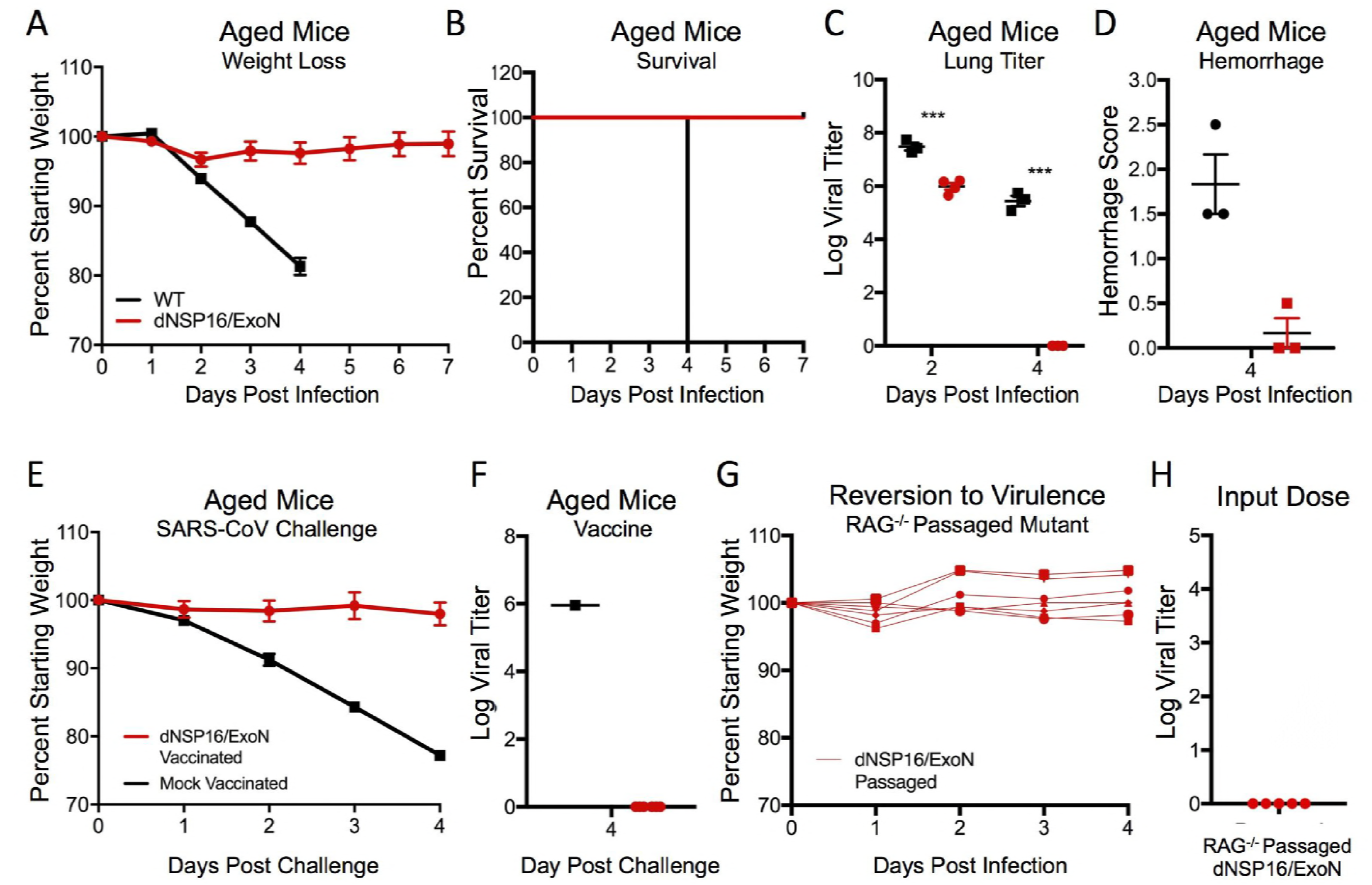
SARS-CoV dNSP16/ExoN protects aged and is cleared from immune compromised mic. A-D) Twelve-month old BALB/c mice were challenged with 100 PFU of WT SARS-CoV MA15 (black), dNSP16/ExoN (red) and examined for A) weight loss, B) survival, C) lung viral titer, and D) lung hemorrhage over the time course. E-F) Twelve-month old BALB/c mice were vaccinated with 100 PFU of SARS-CoV dNSP16/ExoN (red) or mock (PBS, black), monitored for 28 days, and subsequently challenged with 10^5^ PFU of WT SARS-CoV MA15. Mice were examined over a four day time course for E) weight loss and F) lung viral titer. H) Five RAG^-/-^ mice were infected with 10^4^ PFU of SARS-CoV dNSP16/ExoN, monitored, euthanized, and lung tissues harvested after 30 days. Clarified homogenates were then inoculated into 10-week old BALB/c mice and monitored for G) weight loss and H) input titer. P-values based on Student T-test and are marked as indicated: * < 0.05, * * < 0.01, * * * < 0.001.

### SARS-CoV dNSP16/ExoN cleared from immune compromised mice

Having addressed both heterologous challenge and efficacy in aged models, we next examined the dNSP16/ExoN mutation for reversion potential. Similar to the original dNSP16 mutant, dNSP16/ExoN showed no significant disease in RAG^-/-^ mice during acute infection. After 30 days, mice were euthanized, lung tissue harvested, homogenized, clarified, and used to infect 10-week old BALB/c mice. Contrasting dNSP16 (**Fig. 4**), reinfection from lung homogenate of RAG^-/-^ mice infected with the double mutant failed to induce any significant disease in terms of weight loss or lethality upon acute infection (**Fig. 7G**). Examination of input viral titer revealed the absence of detectable virus in all five RAG^-/-^ mice infected with the dNSP16/ExoN mutant (**Fig. 7H**). The results indicate that following *in vivo* replication for 30 days, the dNSP16/ExoN mutant failed to revert to a virulent form. In addition, the results suggest that the double mutant could be cleared in immune compromised mice without the need for traditional adaptive immunity. Together, the data argue that the dNSP16/ExoN mutant is unlikely to revert even in immune compromised models and fulfills a key consideration for a live-attenuated coronavirus platform.

## Discussion

The threat of zoonotic coronaviruses stems not only from their potential for emergence, but also from the lack of effective strategies to combat epidemic coronavirus disease. While both SARS-CoV and MERS-CoV have been primarily limited through effective public health measures, future emergence events may not be as easily or quickly addressed. With this in mind, we evaluated the use of 2’O MTase mutants as a live-attenuated vaccine platform to rapidly respond to current and future CoV strains. Our studies illustrated that despite success in standard young mouse models, the CoV NSP16 mutation alone carried significant risk for pathogenesis in aged animals and had the potential for reversion to virulence. However, pairing the NSP16 mutation with a second attenuating mutation provided a SARS-CoV vaccine capable of protecting against homologous and heterologous challenge, efficacy in aged mice, and no evidence for virus reversion to virulence. Together, the study highlights the potential for NSP16 attenuation as a major feature of future live-attenuated coronavirus vaccine approaches.

Similar to the ExoN and E proteins, NSP16 is conserved across the entire coronavirus family increasing its appeal as an attenuation target. While 2’O MTase activity impacts the immune response and pathogenesis, it is not explicitly required for viral replication as demonstrated by the lack of *in vitro* attenuation. In contrast, both ExoN and E protein mutants have replication defects *in vitro* and have reduced overall yields during virus production. Importantly, this replication attenuation potentially contributes to the difficulty in recovering ExoN and severe attenuation of E protein mutants in the context of MERS-CoV infection. However, the absence of replication defects for NSP16 mutants offers both utility and risk. While capable of inducing robust immunity, even to heterologous challenge, the NSP16 mutant virus retains virulence in aged animals at high doses. Importantly, the augmented replication potentially contributes to its ability to revert to virulence in immunocompromised models. Together, the results suggest that the robust immunity induced by NSP16 mutants in young mice comes at the cost of higher virulence and the potential for reversion, thus limiting its use to only dire situations.

Given the risk associated with NSP16 mutation as a single target attenuation platform, we evaluated ways to increase safety without sacrificing vaccine efficacy. We first considered further targeting the conserved KDKE motif, critical to 2’O MTase function. However, mutations of the SARS NSP16 (K46A, K130A) individually resulted in replication attenuation, potential compensatory mutations, and concerns for mutant viability when used in combination (9). In addition, despite the D130A mutation representing only a two-nucleotide mutation (20, 22), none of the RAG^-/-^ passaged viruses that caused disease had reverted to the original residue. The results indicate that the SARS-CoV dNSP16 mutant replicated sufficiently to permit adaptation at other sites and can restore virulence. Therefore, we instead focused on pairing the NSP16 mutant with another SARS-CoV attenuating mutation in ExoN. The combined double mutant had similar replication deficits as the singly targeted ExoN mutant, but now featured both IFN/IFIT1 and fidelity based attenuation as well. Importantly, the double mutant vaccine protected against heterologous challenge, proved effective in aged models, and showed no evidence of reversion in immune compromised models. Together, the results indicate that targeting NSP16 in combination with another attenuating feature is a viable approach to improve live-attenuated vaccine safety.

In pursuing vaccine platforms for coronaviruses, our data argues a multiple-mechanism, attenuation approach may be the fastest route to an effective, safe vaccine in the context of an outbreak. In this work, the absence of replication deficits permitted pairing of the NSP16 mutation with the less fit SARS ExoN mutant; alternatively, a SARS E protein mutant could also have been selected and has previously been shown to work in combination with another attenuating mutation in SARS NSP1 (17). However, the appeal of targeting NSP16 revolves around the lack of replication attenuation, complete conservation of the required KDKE motif, and the vaccine protection offered by a viable MERS-CoV NSP16 mutant. These factors, absent from both ExoN and E protein platforms, make targeting NSP16 a priority approach for live-attenuated vaccines. Importantly, translation of vaccine efficacy to heterologous challenge, vulnerable aged populations, and lack of reversion to virulence distinguish the NSP16 combination platform from other approaches tried to date.

With the emergence and ongoing outbreak of MERS-CoV in the Middle East, renewed efforts have been placed on effective vaccination approaches (13). Proof of efficacy against MERS-CoV has been explored for several platforms including live-attenuated viruses, DNA plasmids, viral based vectors, nanoparticles, and recombinant protein strategies (11); some of these approaches have even advanced to phase I trials in humans (26). Yet, despite initial successes, a truly effective MERS-CoV vaccine must also consider vulnerable aging populations and the threat from heterologous challenge. Absent surveying these important factors, MERS-CoV vaccine approaches may bear similar failures that beset SARS-CoV vaccines developed a decade earlier (14, 15). Importantly, our studies indicate that a live-attenuated SARS-CoV vaccine incorporating NSP16 and an additional attenuating mutation overcomes these issues and provides protection to vulnerable populations with limited risk for reversion. In pursuit of a platform to rapidly respond to an emerging threat, combination live-attenuated vaccines employing NSP16 mutations may offer the safest path forward.

## Methods & Materials

### Cells & Viruses

Wild-type, mutant, and mouse adapted SARS-CoV were previously described (27, 28) and cultured on Vero 81 cells, grown in DMEM or OptiMEM (Gibco, CA) and 5% Fetal Clone Serum (Hyclone, South Logan, UT) along with antibiotic/antimycotic (Gibco, Carlsbad, CA). Growth curves in Vero cells were performed as previously described with multiple samples (n ≥ 3) examined at each time point (9, 31). Briefly, cells were washed with PBS, and inoculated with virus or mock diluted in PBS for 40 minutes at 37 °C. Following inoculation, cells were washed 3 times, and fresh media added to signify time 0 hr. Samples were harvested at the described time points. All virus cultivation was performed in a BSL3 laboratory with redundant fans in Biosafety Cabinets as described previously by our group (32, 33). All personnel wore Powdered Air Purifying Respirators (3M breathe easy) with Tyvek suits, aprons, booties and were double-gloved.

### Construction of Wild-type and Mutant NSP16, NSP16/ExoN Viruses

Both wild-type and mutant viruses were derived from either SARS-CoV Urbani or corresponding mouse adapted (MA15) infectious clone as previously described (25, 29). For NSP16/ExoN mutant construction, we utilized the constructs previously generated for individual NSP16 (19) and NSP14 ExoN (16) mutants located on different fragments (D & E respectively). Thereafter, plasmids containing wild-type and mutant SARS-CoV genome fragments were amplified, excised, ligated, and purified. *In vitro* transcription reactions were then preformed to synthesize full-length genomic RNA, which was transfected into Vero E6 cells. The media from transfected cells were harvested and served as seed stocks for subsequent experiments. Viral mutants were confirmed by sequence analysis prior to use. Synthetic construction of mutants of NSP16 and NSP14 were approved by the University of North Carolina Institutional Biosafety Committee.

### RNA isolation, microarray processing and identification of differential expression

RNA isolation and microarray processing, quality control, and normalization from whole lung homogenate was carried out as previously described (30). Differential expression (DE) was determined by comparing virus-infected replicates to time-matched mock replicates. Criteria for DE in determining the consensus ISG list were an absolute Log_2_ fold change of >1.5 and a false discovery rate (FDR)-adjusted *P* value of <0.05 for a given time point.

### Clustering and Functional Enrichment

Genes identified as differentially expressed were used to generate clustered expression heat maps. Hierarchical clustering (using Euclidean distance and complete linkage clustering) was used to cluster gene expression according to behavior across experimental conditions. Mouse GO terms (www.geneontology.org) and the EASE-adjusted fisher exact test (31) were used to determine functional enrichment results for the genes in each cluster. Functional enrichment output was manually summarized for each cluster. PCA plot and cluster diagram were generated using the prcomp() function in the R base package, and the gplots R package, respectively.

### Ethics Statement

This study was carried out in accordance with the recommendations for care and use of animals by the Office of Laboratory Animal Welfare (OLAW), National Institutes of Health. The Institutional Animal Care and Use Committee (IACUC) of The University of North Carolina at Chapel Hill (UNC, Permit Number A-3410-01) approved the animal study protocol (IACUC Protocol #15-009 and 13-072) followed in this manuscript.

### Mouse Infections

10-week old mice and 12-month old mice were anaesthetized with ketamine and Xylazine (as per IACUC, UNC guidelines) and intranasally inoculated with a 50 μl volume containing 10^2^ to 10^5^ plaque forming units (PFU) of SARS-CoV MA15, SARS-CoV dNSP16, SARS-CoV dNSP16/ExoN virus, or PBS mock as indicated in the figure legends. For vaccination experiments, mice were infected with vaccination dose of SARS-CoV dNSP16 or SARS-CoV dNSP16/ExoN as described above, monitored for clinical symptoms for 7 days, and then challenged 4 weeks post vaccination with 10^5^ PFU SARS-CoV MA15. Infected animals were monitored for weight loss, morbidity, clinical signs of disease, and lung titers were determined as described previously (6). Young and aged female BALB/c mice were purchased from Envigo/Harlan Labs; female RAG^-/-^ mice were obtained from The Jackson Laboratory (Bar Harbor, Maine).

### Virus Neutralization Assays

Plaque reduction neutralization titer assays were performed with previously characterized antibodies against SARS-CoV as previously described (32). Briefly, neutralizing antibodies or serum were serially diluted 2-fold and incubated with 100 PFU of the different virus strains for 1 h at 37 °C. The virus and antibodies were then added to a 6-well plate with 5 × 10^5^ Vero E6 cells/well with n ≥ 2. After an 1-h incubation at 37 °C, cells were overlaid with 3 ml of 0.8% agarose in media. Plates were incubated for 2 days at 37 °C and then stained with neutral red for 3 h, and plaques were counted. The percentage of plaque reduction was calculated as [1 - (no. of plaques with antibody/no. of plaques without antibody)] × 100.

### Data Dissemination

Raw microarray data for these studies were deposited in publicly available databases in the National Center for Biotechnology Information’s (NCBI) Gene Expression Omnibus (3) and are accessible through GEO Series: GSE49263. (http://www.ncbi.nlm.nih.gov/geo/query/acc.cgi?acc=GSE49263).

## Acknowledgements

Research was supported by grants from NIAID of the NIH (U19AI100625, U19AI106772, and AI108197 to RSB; R00AG049092 to VDM). The content is solely the responsibility of the authors and does not necessarily represent the official views of the NIH. PNNL is operated by Battelle Memorial Institute for the DOE under contract number DE-AC05-76RLO1830.

## References

1. Perlman S, Netland J. 2009. Coronaviruses post-SARS: update on replication and pathogenesis. Nat Rev Microbiol 7:439–450.

2. Cunningham AA, Daszak P, Wood JLN. 2017. One Health, emerging infectious diseases and wildlife: two decades of progress? Philos Trans R Soc Lond B Biol Sci 372.

3. Zaki AM, van Boheemen S, Bestebroer TM, Osterhaus AD, Fouchier RA. 2012. Isolation of a novel coronavirus from a man with pneumonia in Saudi Arabia. N Engl J Med 367:1814–1820.

4. Menachery VD, Graham RL, Baric RS. 2017. Jumping species-a mechanism for coronavirus persistence and survival. Curr Opin Virol 23:1–7.

5. Fehr AR, Channappanavar R, Perlman S. 2017. Middle East Respiratory Syndrome: Emergence of a Pathogenic Human Coronavirus. Annu Rev Med 68:387–399.

6. Stockman LJ, Bellamy R, Garner P. 2006. SARS: systematic review of treatment effects. PLoS Med 3:e343.

7. Menachery VD, Yount BL, Jr., Debbink K, Agnihothram S, Gralinski LE, Plante JA, Graham RL, Scobey T, Ge XY, Donaldson EF, Randell SH, Lanzavecchia A, Marasco WA, Shi ZL, Baric RS. 2015. A SARS-like cluster of circulating bat coronaviruses shows potential for human emergence. Nat Med 21:1508–1513.

8. Kilianski A, Baker SC. 2014. Cell-based antiviral screening against coronaviruses: developing virus-specific and broad-spectrum inhibitors. Antiviral Res 101:105112.

9. Sheahan TP, Sims AC, Graham RL, Menachery VD, Gralinski LE, Case JB, Leist SR, Pyrc K, Feng JY, Trantcheva I, Bannister R, Park Y, Babusis D, Clarke MO, Mackman RL, Spahn JE, Palmiotti CA, Siegel D, Ray AS, Cihlar T, Jordan R, Denison MR, Baric RS. 2017. Broad-spectrum antiviral GS-5734 inhibits both epidemic and zoonotic coronaviruses. Sci Transl Med 9.

10. Graham RL, Donaldson EF, Baric RS. 2013. A decade after SARS: strategies for controlling emerging coronaviruses. Nat Rev Microbiol 11:836–848.

11. Zumla A, Chan JF, Azhar EI, Hui DS, Yuen KY. 2016. Coronaviruses - drug discovery and therapeutic options. Nat Rev Drug Discov 15:327–347.

12. Menachery VD, Mitchell HD, Cockrell AS, Gralinski LE, Yount BL, Jr., Graham RL, McAnarney ET, Douglas MG, Scobey T, Beall A, Dinnon K, 3rd, Kocher JF, Hale AE, Stratton KG, Waters KM, Baric RS. 2017. MERS-CoV Accessory ORFs Play Key Role for Infection and Pathogenesis. MBio 8.

13. Chafekar A, Fielding BC. 2018. MERS-CoV: Understanding the Latest Human Coronavirus Threat. Viruses 10.

14. Bolles M, Deming D, Long K, Agnihothram S, Whitmore A, Ferris M,Funkhouser W, Gralinski L, Totura A, Heise M, Baric RS. 2011. A doubleinactivated severe acute respiratory syndrome coronavirus vaccine provides incomplete protection in mice and induces increased eosinophilic proinflammatory pulmonary response upon challenge. J Virol 85: 12201–12215.

15. Deming D, Sheahan T, Heise M, Yount B, Davis N, Sims A, Suthar M, Harkema J, Whitmore A, Pickles R, West A, Donaldson E, Curtis K, Johnston R, Baric R. 2006. Vaccine efficacy in senescent mice challenged with recombinant SARS-CoV bearing epidemic and zoonotic spike variants. PLoS Med 3: e525.

16. Graham RL, Becker MM, Eckerle LD, Bolles M, Denison MR, Baric RS. 2012. A live, impaired-fidelity coronavirus vaccine protects in an aged, immunocompromised mouse model of lethal disease. Nat Med 18: 1820–1826.

17. Jimenez-Guardeno JM, Regla-Nava JA, Nieto-Torres JL, DeDiego ML, Castano-Rodriguez C, Fernandez-Delgado R, Perlman S, Enjuanes L. 2015. Identification of the Mechanisms Causing Reversion to Virulence in an Attenuated SARS-CoV for the Design of a Genetically Stable Vaccine. PLoS Pathog 11: e1005215.

18. Zust R, Cervantes-Barragan L, Habjan M, Maier R, Neuman BW, Ziebuhr J, Szretter KJ, Baker SC, Barchet W, Diamond MS, Siddell SG, Ludewig B, Thiel V. 2011. Ribose 2’-O-methylation provides a molecular signature for the distinction of self and non-self mRNA dependent on the RNA sensor Mda5. Nat Immunol 12: 137-143.

19. Menachery VD, Yount BL, Jr., Josset L, Gralinski LE, Scobey T, Agnihothram S, Katze MG, Baric RS. 2014. Attenuation and restoration of severe acute respiratory syndrome coronavirus mutant lacking 2’-o-methyltransferase activity. J Virol 88:4251–4264.

20. Menachery VD, Gralinski LE, Mitchell HD, Dinnon KH, 3rd, Leist SR, Yount BL, Jr., Graham RL, McAnarney ET, Stratton KG, Cockrell AS, Debbink K, Sims AC, Waters KM, Baric RS. 2017. Middle East Respiratory Syndrome Coronavirus Nonstructural Protein 16 Is Necessary for Interferon Resistance and Viral Pathogenesis. mSphere 2.

21. DeDiego ML, Nieto-Torres JL, Jimenez-Guardeno JM, Regla-Nava JA, Castano-Rodriguez C, Fernandez-Delgado R, Usera F, Enjuanes L. 2014. Coronavirus virulence genes with main focus on SARS-CoV envelope gene. Virus Res 194:124-137.

22. Menachery VD, Debbink K, Baric RS. 2014. Coronavirus non-structural protein 16: evasion, attenuation, and possible treatments. Virus Res 194:191–199.

23. Roberts A, Paddock C, Vogel L, Butler E, Zaki S, Subbarao K. 2005. Aged BALB/c mice as a model for increased severity of severe acute respiratory syndrome in elderly humans. J Virol 79:5833–5838.

24. Sheahan T, Whitmore A, Long K, Ferris M, Rockx B, Funkhouser W, Donaldson E, Gralinski L, Collier M, Heise M, Davis N, Johnston R, Baric RS. 2011. Successful vaccination strategies that protect aged mice from lethal challenge from influenza virus and heterologous severe acute respiratory syndrome coronavirus. J Virol 85:217–230.

25. Roberts A, Deming D, Paddock CD, Cheng A, Yount B, Vogel L, Herman BD, Sheahan T, Heise M, Genrich GL, Zaki SR, Baric R, Subbarao K. 2007. A mouse-adapted SARS-coronavirus causes disease and mortality in BALB/c mice. PLoS Pathog 3:e5.

26. Modjarrad K. 2016. MERS-CoV vaccine candidates in development: The current landscape. Vaccine 34:2982–2987.

27. Scobey T, Yount BL, Sims AC, Donaldson EF, Agnihothram SS, Menachery VD, Graham RL, Swanstrom J, Bove PF, Kim JD, Grego S, Randell SH, Baric RS. 2013. Reverse genetics with a full-length infectious cDNA of the Middle East respiratory syndrome coronavirus. Proc Natl Acad Sci U S A 110:16157–16162.

28. Cockrell A YB, Scobey T, Jensen K, Douglas M, Beall A, Tang X-C, Marasco WA, Heise MT, Baric RS 2016. A Mouse Model for MERS Coronavirus Induced Acute Respiratory Distress Syndrome. Nature Microbiology In Press.

29. Yount B, Curtis KM, Fritz EA, Hensley LE, Jahrling PB, Prentice E, Denison MR, Geisbert TW, Baric RS. 2003. Reverse genetics with a full-length infectious cDNA of severe acute respiratory syndrome coronavirus. Proc Natl Acad Sci U S A 100:12995–13000.

30. Gralinski LE, Bankhead A, 3rd, Jeng S, Menachery VD, Proll S, Belisle SE, Matzke M, Webb-Robertson BJ, Luna ML, Shukla AK, Ferris MT, Bolles M, Chang J, Aicher L, Waters KM, Smith RD, Metz TO, Law GL, Katze MG, McWeeney S, Baric RS. 2013. Mechanisms of severe acute respiratory syndrome coronavirus-induced acute lung injury. MBio 4.

31. Hosack DA, Dennis G, Jr., Sherman BT, Lane HC, Lempicki RA. 2003. Identifying biological themes within lists of genes with EASE. Genome Biol 4:R70.

32. Menachery VD, Yount BL, Jr., Sims AC, Debbink K, Agnihothram SS, Gralinski LE, Graham RL, Scobey T, Plante JA, Royal SR, Swanstrom J, Sheahan TP,Pickles RJ, Corti D, Randell SH, Lanzavecchia A, Marasco WA, Baric RS. 2016. SARS-like WIV1-CoV poised for human emergence. Proc Natl Acad Sci U S A 113:3048–3053.

33. Sims AC, Tilton SC, Menachery VD, Gralinski LE, Schafer A, Matzke MM, Webb-Robertson BJ, Chang J, Luna ML, Long CE, Shukla AK, Bankhead AR, 3rd, Burkett SE, Zornetzer G, Tseng CT, Metz TO, Pickles R, McWeeney S, Smith RD, Katze MG, Waters KM, Baric RS. 2013. Release of severe acute respiratory syndrome coronavirus nuclear import block enhances host transcription in human lung cells. J Virol 87:3885–3902.

